# A novel viral protein translation mechanism reveals mitochondria as a target for antiviral drug development

**DOI:** 10.1101/2020.10.19.344713

**Authors:** Zhenguo Cheng, Danhua Zhang, Jingfei Chen, Yifan Wu, XiaoWen Liu, Lingling Si, Zhe Zhang, Na Zhang, Zhongxian Zhang, Wei Liu, Hong Liu, Lirong Zhang, Lijie Song, Louisa S Chard Dunmall, Jianzeng Dong, Nicholas R Lemoine, Yaohe Wang

## Abstract

The ongoing Severe Acute Respiratory Syndrome coronavirus 2 (SARS-CoV-2) pandemic has acutely highlighted the need to identify new treatment strategies for viral infections. Here we present a pivotal molecular mechanism of viral protein translation that relies on the mitochondrial translation machinery. We found that rare codons such as Leu-TTA are highly enriched in many viruses, including SARS-CoV-2, and these codons are essential for the regulation of viral protein expression. SARS-CoV-2 controls the translation of its spike gene by hijacking host mitochondria through 5’ leader and 3’UTR sequences that contain mitochondrial localization signals and activate the EGR1 pathway. Mitochondrial-targeted drugs such as lonidamine and polydatin significantly repress rare codon-driven gene expression and viral replication. This study identifies an unreported viral protein translation mechanism and opens up a novel avenue for developing antiviral drugs.

**One Sentence Summary:** Mitochondria are a potential target for antiviral therapy

## Main Text

Severe acute respiratory syndrome coronavirus 2 (SARS-CoV-2), the etiological agent responsible for the ongoing pandemic of COVID-19, has killed more than one million people in around 200 countries since its identification in December 2019. There are currently three main categories of anti-viral drug being assessed for use in COVID-19 patients: viral replication inhibitors that target viral proteins; drugs that block host cell proteins involved in viral infection, and reagents that balance host immune responses to infection(*1, 2*). Unfortunately, effective antiviral drugs are still lacking. Recently the interaction network of viral proteins(*3*), the global landscape of host gene expression(*4, 5*) and host and viral phosphoproteomics(*6*) after SARS-CoV-2 infection have been reported and this information may accelerate the screening of potential antiviral agents. However, the detailed mechanisms by which viral genes interact with host cell intracellular compartments remain largely unknown. Defining these mechanisms is important not only for understanding viral pathogenesis but also for designing new antiviral therapeutics.

Given the degeneracy of the genetic code, all amino acids are coded by 2-6 synonymous codons that are not used with equal frequency. Codon usage bias, the preference for certain synonymous codons for each amino acid, is a universal feature of all genomes and plays an important role in determination of gene expression levels. A positive correlation between codon usage bias and tRNA expression allows more efficient translation of genes enriched in optimal codons compared to those enriched in rare codons(*7*). The presence of optimal codons has been associated with an enhanced rate of translation elongation, the control of ribosome traffic on the mRNA and regulation of proper co-translational protein folding(*8*). Understanding the codon usage bias in different organisms has implications for future therapeutic design.

Most viruses have compact genomes lacking transfer RNA (tRNA) coding sequences, thus translation of viral proteins is completely reliant on host cell translation machinery(*9*). Recent studies have demonstrated that viruses can effectively manipulate host tRNAs to better match viral codon usage, presenting a promising target for restriction of viral replication(*10, 11*).

SARS-CoV-2 is a single-stranded, positive-sense group β coronavirus, with a genome of around 30kbp (*12*). The virus genome contains 10 open-reading frames (ORFs) that encode four structural proteins: Spike (S), Membrane (M), Envelope (E) and the nucleocapsid (N) proteins, and six non-structural proteins ORF1ab, ORF3, ORF6, ORF7, ORF8 and ORF10 (*13*). We found that the native sequence of the SARS-CoV-2 spike (S) gene expressed protein at an extremely low level after transfection of recombinant plasmid into human cells, but expression was significantly improved after codon optimization of the S protein gene (Fig. 1A)(*14*). To better understand the protein expression pattern of SARS-CoV-2, protein abundance of the four structural proteins (S, E, M and N) and four accessory proteins (ORF3, ORF6, ORF7 and ORF8) was evaluated and we found that the protein expression of M, N, ORF3 and ORF7 could be readily detected in human cells after the transfection of a plasmid expressing native gene sequences, while the abundance of S, E, ORF6 and ORF8 were low, with no observed positive band when the native gene sequences were used (Fig. 1B). These results were corroborated using immunofluorescence, in which the viral ORFs were fused with RFP using an IRES motif (*15*) (Fig. S1). These findings indicate that several key genes of SARS-CoV-2 show codon usage bias and dissection of the underlying mechanisms responsible may provide a new direction for targeting of antiviral drugs.

**Fig.1.**
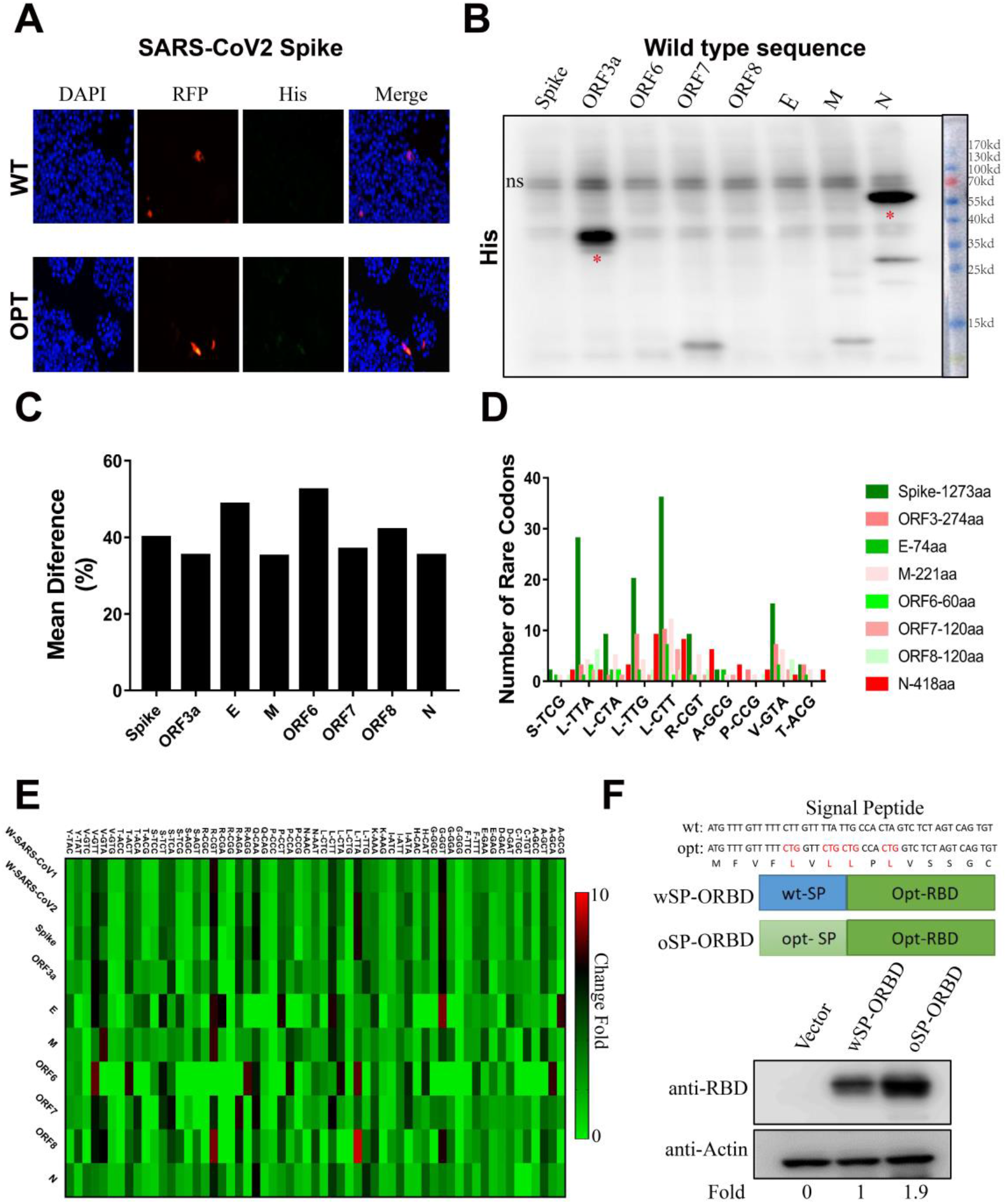
Rare codon bias can prevent translation of SARS-CoV-2 derived sequences. (A) Protein expression of plasmids expressing SARS-CoV-2 S using the original or codon-optimized sequence after transfection into human A549 cells. Expression was evaluated using confocal microscopy, and an IRES-RFP sequence was included after the S open reading frame (ORF) for visualisation. (B) Western Blot assay was performed to detect the protein abundance of SARS-CoV-2-derived sequences including the four structural proteins (spike (S), envelope (E), membrane (M) and nucleocapsid (N)) and four accessory proteins (ORF3, ORF6, ORF7 and ORF8). The abbreviation ‘ns’ indicates non-specific. (C) Codon usage variability of selected SARS-CoV-2 ORFs was analysed with GCUA software using the human standard codon table as reference, and the figure was drawn using GraphPad Prism. (D) The number of rare codons (fraction <0.15 and frequency/thousands <15) in the eight viral genes sequences. (E) Relative change of codon usage (not containing codon “ATG”, “TGG”, “TAA”, “TAG” and “TGA”) in SARS-CoV1, SARS-CoV2 and eight viral genes coding sequence using the human standard codon table as control. (F) The effect of rare codon “Leu” on protein expression was assessed by comparing expression of wild-type or optimized signal peptide of spike (signal peptide of spike has several “Leu”, the rare codons encoding “Leu” were replaced with the most commonly used codon CTG (in red colour)), Both sequences included a codon-optimized RBD domain (RBD could be replaced by other protein).

To confirm SARS-CoV-2 codon usage, the rare codon usage preference among SARS-CoV-2 and humans were analyzed using the GCUA tool. As expected, viral proteins with low-abundance (S, E, ORF6, ORF8) were more highly enriched in rare codons than highly expressed proteins (M, N, ORF3, ORF7) (Fig. 1C). Several rare codons, especially those encoding Leucine (Leu), were present in higher numbers and proportion in low-abundance proteins (Fig. 1D & 1E). Sequence analysis revealed that the 5’ signal peptide of the S protein possesses several rare Leu codons. To validate the effect of these rare codons on S protein expression, the wild-type signal peptide or codon-optimized signal peptide, in which the four rare codons were replaced with a human optimized codon Leu-CTG, were fused in frame with the codon-optimized receptor binding domain (RBD) of S, which is responsible for interactions with the angiotensin converting enzyme 2 (ACE2) receptor to mediate viral cell entry. RBD domain protein expression was significantly improved by codon optimisation (Fig. 1F), suggesting that the presence of the tRNAs required for decoding low-usage Leu codons may be crucial for translation of SARS-CoV-2 S protein and as such, a potential intervention target.

Considering the importance of tRNAs in translation, genome analysis of TRL-TAA (the tRNA specific for Leu, expressing the TAA anticodon) revealed a strong signal of transcriptional factor (TF) binding and histone modification around TRL-TAA (Fig. 2A). Potential virus-associated TFs interacting with TRL-TAA promoter were predicted with JASPAR software (Tables S1), and 25 elevated TFs were identified (Fig. 2B & C), indicating these TFs may affect the translation of viral genes enriched in rare codons. Subsequently, the top candidate, early growth response 1 (EGR1), was evaluated for a role in rare codon usage. Despite previous studies demonstrating that host EGR1 induced by Vaccinia virus or cytomegalovirus infection enhances virus replication and infectivity (*16, 17*), EGR1 overexpression in HEK293 cells was only able to slightly increase S protein expression with wild type sequence (Fig. 2D). To further investigate the potential accessory TFs of SARS-CoV-2 infection, signalling pathway enrichment was performed to identify the relationship of those genes with increased expression in SARS-CoV-2-infected cells (GSE147507). Data showed these genes were mainly clustered in pathways including TNF signalling, cytokine production, ATF2 pathway and inflammatory response (Fig. 2E). However, overexpression of activating transcription factor 2 (ATF2) had no significant effect on S expression (Fig. 2D),despite the promoter of TRL-TAA also contains potential binding sites for ATF2 (Tables S1). Interestingly, the data of human single-cell transcriptome analysis experiments revealed that most of these TFs were enriched in ACE2- and TMPRSS2-positive cells (*18*), especially in ciliated cells and pneumocytes, SARS-CoV-2 target cells (Fig. 2F) (*19, 20*).

**Fig.2.**
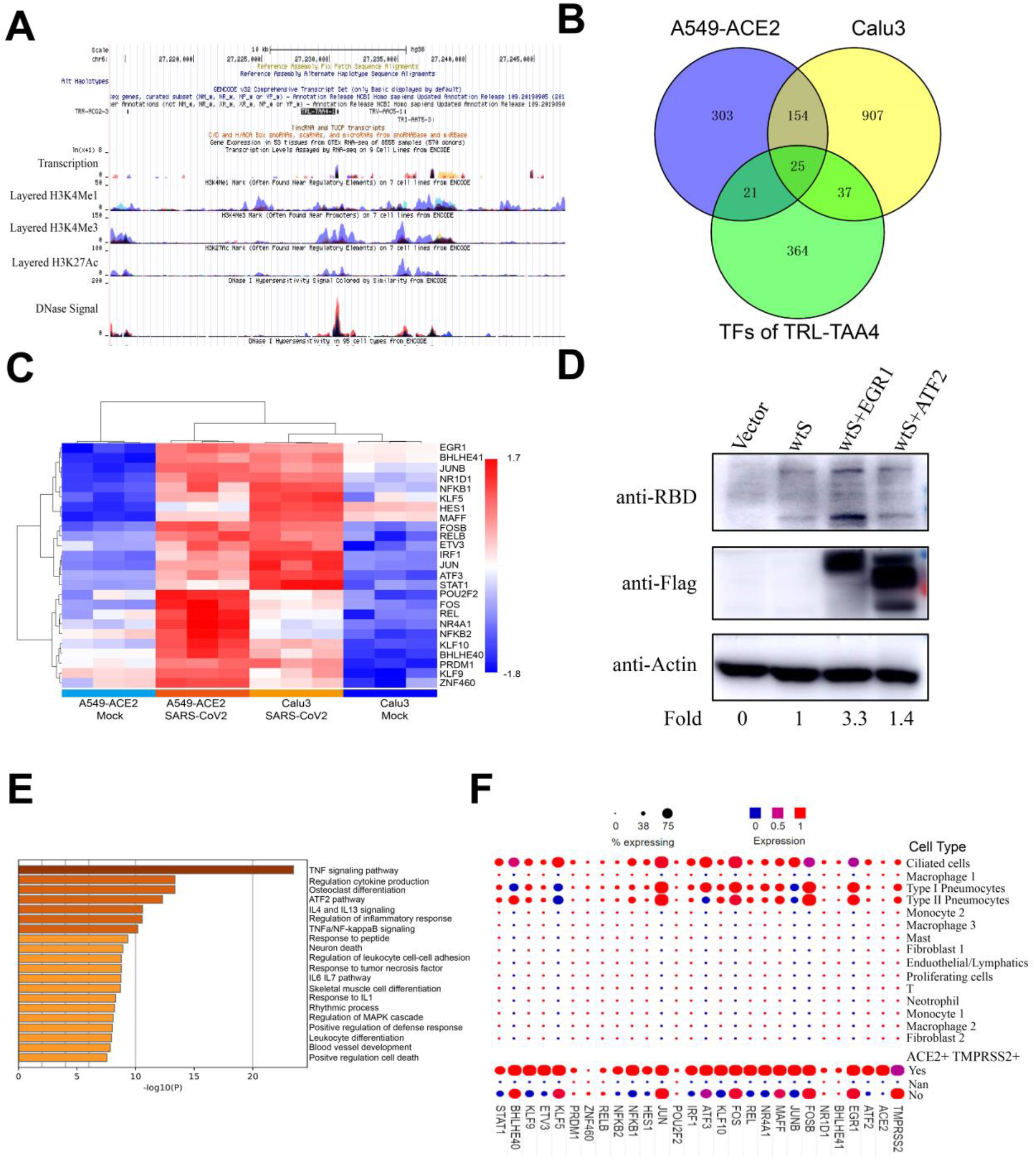
The possible effect of Transcription factors binding Leucine-specific tRNAs in viral protein translation. (A) Genomic characteristics including histone modification and transcription factor binding status of tRNA TRL-TAA4-1 which matched with rare codon Leu-TTA was analysed using the UCSC database. (B) Overlap analysis of potential TRL-TAA regulatory transcription factors (TFs) and genes that increased in SARS-CoV-2 infection in A549-ACE2 and Calu3 cells using Venny2.1. (C) Heat map demonstrating the expression of 25 TFs in SARS-CoV-2-infected cell lines created using R studio. (D) Western Blot assay to assess the protein level of S after co-transfection with EGR1 or ATF2 in HEK-293 cells. (E). Differentially expressed genes after SARS-CoV-2 infection (GSE147507) were analyzed with R Studio. Pathway enrichment of highly expressed genes was mapped using Metascape (F) Distribution of genes including ACE2, TMPRSS2, ATF2 and identified TFs in human lung cells was created using public single cell sequence data(*18*).

Given that after natural infection, intrinsic S protein expression levels in virus-infected cells are high(*5*), we hypothesized that there may be other mechanisms that regulate the protein expression of S and other genes with rare codons. To confirm this, enrichment analysis was performed using genes reported to interact with S (*3*). As shown in Fig. 3A, these proteins were mainly clustered to protein folding, mRNA splicing and protein translation functions, in particular proteins involved in the process of mitochondrial gene translation.

**Fig.3.**
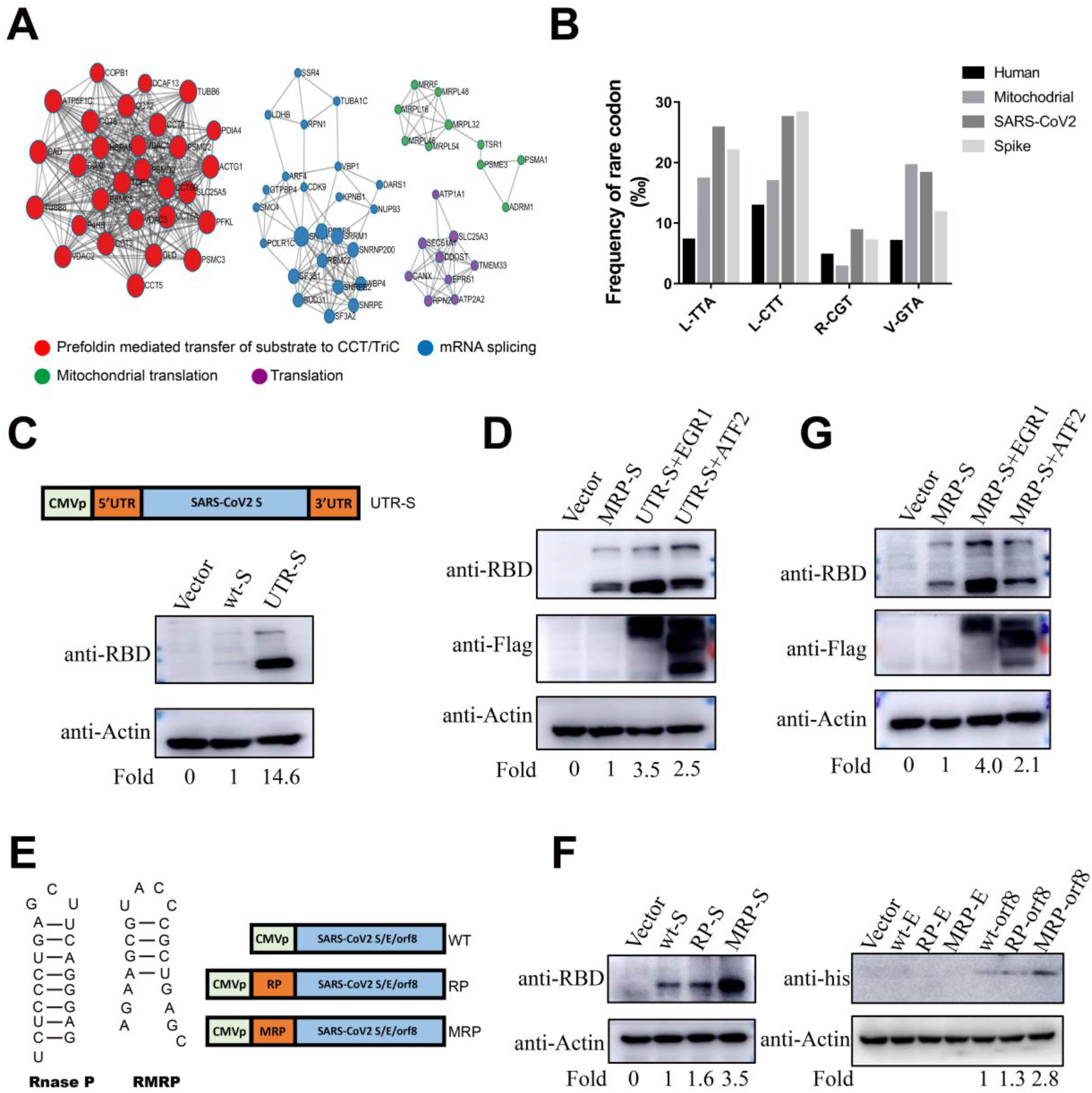
Mitochondrial localization is critical for the translation of SARS-CoV-2 S protein. (A) The interaction network of proteins that bound with S was analyzed using Metascape. (B) The frequency of rare codons in SARS-CoV-2, S protein, human nuclear genome and the human mitochondrial genome using the Codon Usage Database. (C) The protein expression of S flanked by SARS-CoV-2 5’- and 3’-UTR sequences in HEK-293 cell lines. (D) The effect of EGR1 and ATF2 on S protein (plus SARS-CoV-2 5’- and 3’ UTS sequences) expression was analysed using Western blot after transfection into HEK-293 cells. Fold change was determined using Image J software. (E) The skeleton of recombinant plasmids encoding S with or without mitochondrial localisation signals (MLS) RnaseP or RMRP. (F) Protein expression of SARS-CoV-2 derived sequences including S, E and ORF8 and variants expressing MLS as analysed by Western blot after transfection into HEK-293 cells. (G) The effect of EGR1 and ATF2 on wild type spike with RMRP binding motif (MRP).

Eukaryotic cells possess two distinct translation machineries; the cytosolic system that synthesises the majority of proteins within the cell, and the mitochondrial system, specialised for the production of the mitochondrial proteins (*21*). Mitochondria are dynamic organelles that control a wide range of cellular processes, including ATP generation, cellular differentiation, apoptosis and anti-viral immune activation(*22*). The human mitochondrial genome contains only 37 genes, of which two are ribosomal RNAs (rRNAs), 22 tRNAs and 13 protein-coding genes. Mammalian mitochondrial DNA (mtDNA) has neither introns nor protective histones, has a high AT content and as a consequence of the small genome size, employs only a small number of tRNAs which are responsible for decoding a large number of codons (*21*), suggesting that mitochondria show a different codon usage preference from the nuclear genome (*21, 23*). We performed a comparison of codon usage bias and demonstrated that the key rare codon rate in SARS-CoV-2 and S protein were highly similar to the rate seen in mitochondrial genes, and these rates were much higher than found in the human nuclear genome (Fig. 3B). Recently, the SARS-CoV-2 transcriptome revealed that nearly half of SARS-CoV-2 RNAs had 5’ leader sequence (or 5’ untranslated regions, 5’UTR), and both the 5’- and 3’ UTRs of SARS-CoV-2 had potential mitochondrial localization signals(*24*). Dats showed that S protein expression was significantly strengthened in human HEK293 kidney cells by adding the 5’-and 3’UTR sequences to the native S gene in the expression plasmid (Fig. 3C). Moreover, this mitochondrial-dependent protein translation could be further enhanced by overexpression of EGR1 and ATF2 in cells (Fig. 3D), and both proteins have been shown to be related with the function of mitochondria (*25, 26*). To prove further that mitochondrial localization was necessary for effective viral protein translation, two reported mitochondrial RNA targeting signals (Rnase P binding motif, RP and MRP binding motif, MRP) were inserted upstream of the SARS-CoV-2 S, E or ORF8 genes (Fig.3E). The protein level of S and ORF8 with RP or MRP motif were significantly enhanced, although no effect was found regarding expression of E protein with either mitochondrial targeting signal (Fig. 3F). Recently, our results showed that E may play a greater role in viral transcription regulation (data not shown), and this also explained why SARS-CoV without E could still replicate in Vero E6 cells(*27*).

To investigate whether mitochondrial-dependent protein translation was a general principle for the translation of viral proteins, codon bias of 17 viruses including 12 RNA viruses and five DNA viruses were analysed. Impressively, all viruses analysed adopted a codon preference that was more similar to mitochondria (Fig. 4A). Moreover, the fiber capsid protein of adenovirus type 11 (Ad11) which contained rare codons, in particular L-TAA and L-CTT (Fig.4B) and strikingly, modifying the fiber of Ad11 with the MRP motif to enhance mitochondrial localisation was able to increase protein expression compared to expression from the native fiber gene sequence (Fig. 4C). These findings suggest a novel viral protein translation mechanism that relies on manipulation of the mitochondrial translation apparatus.

**Fig.4.**
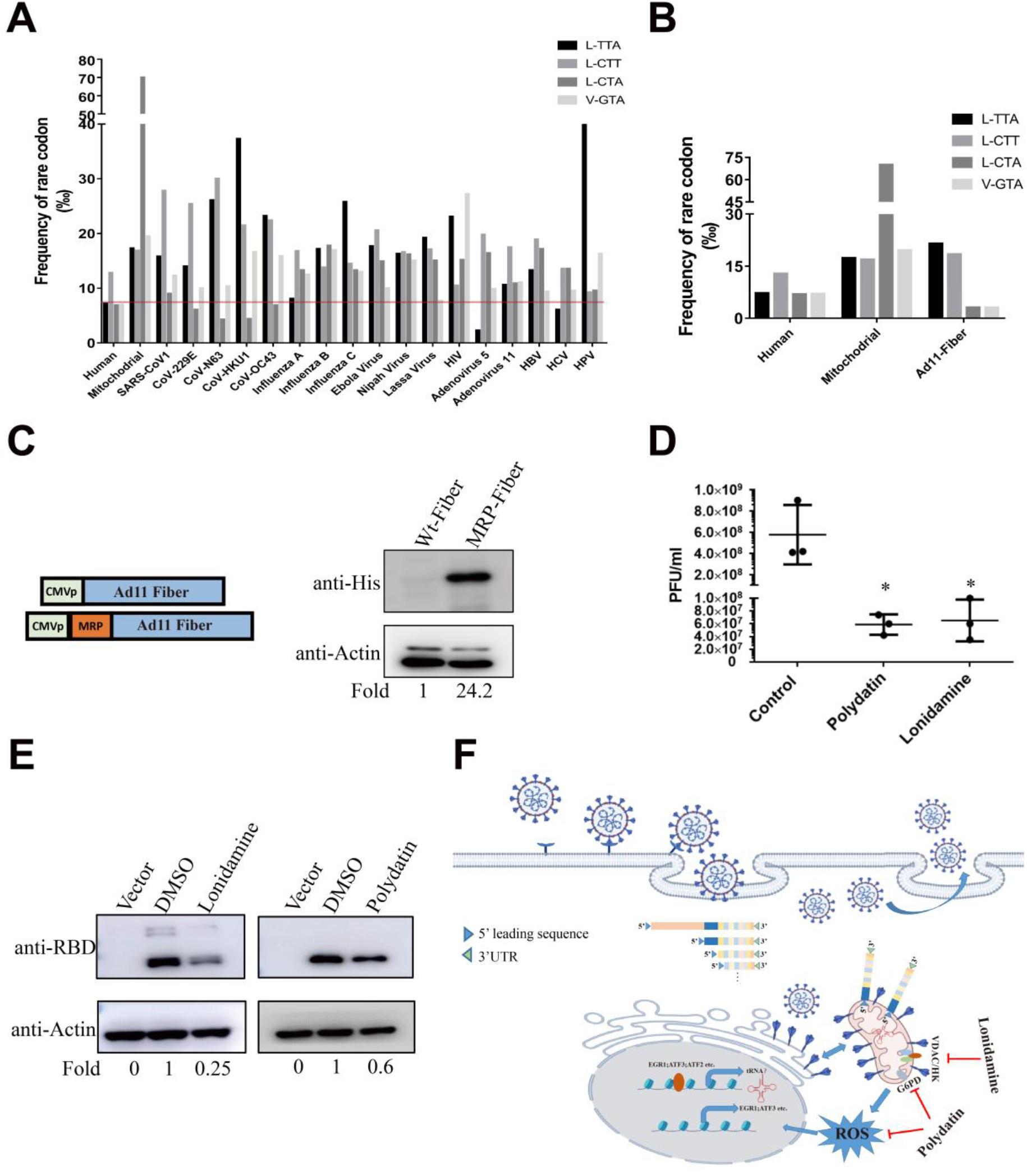
Mitochondrial-targeting drugs have strong potential as anti-viral therapeutic reagents. (A) Codon preferences in 12 RNA viruses and five DNA viruses were downloaded from the Codon Usage Database and the graph was drawn using GraphPad Prism. (B) The codon bias of Ad11 fiber protein was analysed with the Countcodon program in the Codon Usage Database. (C) Protein expression of Ad11 fiber was determined by Western blotting after transfection into HEK-293 cells after transfection with the plasmid containing Ad11 fiber native gene with or without MRP sequences. (D) Human A549 cells were infected with Ad11 virus for 1 hour at MOI=0.5 PFU/cell and further incubated with fresh medium containing 10 μM polydatin or lonidamine for 72 hours. Virus was titrated by TCID assay using HEK293 cells (*, p<0.05). (E) HEK293 cells were transfected with UTR-S constructs and treated with mitochondria-targeted drugs polydatin or lonidamine for 36 hours. Total protein was detected using Western Blot assay. (F) Schematic diagram of the proposed hypothesis regarding the viral hijacking of the mitochondrial translation machinery to support viral infection.

Lonidamine and Polydatin are two well-characterised anti-cancer therapeutics that target mitochondrial pathways to induce cancer cell-death. Treatment with these two drugs significantly repressed Ad11 virus replication (Fig. 4D) and SARS-CoV-2 S protein expression (Fig. 4E) *in vitro*. Interestingly, during the preparation of this manuscript, two Polydatin analogs, Resveratrol and Pterostilbene, both also modulators of mitochondrial function, have been confirmed to have anti-SARS-CoV-2 activity at a time point between the viral entry and release of the virion(*28*). Moreover, another mitochondrion-related drug, 2-deoxy-D-glucose (2DG) was also found to prevent the replication of SARS-COV-2 in Caco-2 cells (*5*). All these findings support the findings presented here regarding the importance of mitochondria for viral protein translation and demonstrate that mitochondria are a promising target for development of antiviral drugs. Both Lonidamine and Polydatin have been tested in human Phase II or III clinical trials for cancer treatment and have demonstrated good safety (*29*). This study suggests that immediate clinical trials for testing their application for SARS-COV-2 and other viral infections are warranted.

In summary, this study has clarified the influence of codon bias on the abundance of rare codon-rich viral proteins such as SARS-CoV-2 S and Ad11 fiber. It has previously been shown that viruses actively interfere with mitochondrial pathways to impede mitochondrial anti-viral signalling mechanisms(*30*). Indeed, SARS-CoV-1 ORF-9b has been demonstrated to localise to mitochondria to manipulate mitochondrial function and suppress anti-viral signalling. The resultant mitochondrial stress also promotes the formation of mitochondrion-derived vesicles that can act as a replication niche for the virus (*31, 32*). Here, for the first time, we disclose that mitochondrial hijacking by viruses and mitochondria localisation of viral RNA are essential for the translation of viral proteins (Fig. 4F). This undoubtedly expands our understanding of the role of mitochondria in viral infections and provides a new insight for the development of antiviral drugs.

## Materials and Methods

### Plasmids

Nucleic acid sequence of four structural proteins including Spike glycoproteins (S), Membrane protein (M), Envelope (E) protein, the nucleocapsid protein (N), and four non-structural proteins including ORF3, ORF6, ORF7, ORF8 derived SARS-CoV-2 were synthesized by Sangon Biotech according to the public sequence provided in the NCBI database (MN908947.3). The genes were digested and cloned into pcDNA3.1A vector to construct the expression plasmids. For recombinant plasmids with IRES-RFP, sequence was amplified from pCMMP-BiP-IRES-RFP (Addgene:#36975) with the following primers IRES-RFP-F: AAGTCGACACGTTACTGGCCGAAGCCG, IRES-RFP-R: AAGGTACCTTAGGCGCCGGTGGAGTGG. In order to construct plasmids with SARS-CoV-2 UTR, we first generated a recombinant vector containing the 5’ leader sequence and 3’UTR, then virus-derived sequences were inserted into the multiple cloning sites available. To obtain recombinant plasmids with mitochondrial localization signal, the oligonucleotide primers of two reported mitochondrial targeting nucleic acid signals (Rnase P-binding motif marked as RP and RMRP binding motif marked as MRP) were synthesized, then the primer dimers were inserted into the 5’ terminal of the genes after denaturation and annealing with the following primer, RP-F: TCTCCCTGAGCTTCAGGGAGTTAATTAAG, RP-R: GATCCTTAATTAACTCCCTGAAGCTCAGGGAGAAT, MRP-F: AGAAGCGTATCCCGCTGAGCTTAATTAAG, MRP-R: GATCCTTAATTAAGCTCAGCGGGATACGCTTCTAT. Human EGR1 clone with flag tag was bought from origene (RC209956), and the human ATF2 expressing clone had been constructed in our previous study(*33*). The plasmid encoding Ad11 fiber was constructed with the following primers, Fiber-F: TCAAGCTTACCGGTGCCACCATGACCAAGAGAGTCCGGCTCAGT, Fiber-R: ACGGATCCTCCGCCACCGCTACCTCCGCCACCGTCGTCTTCTCTGATGTAGTAAAAGGT.

### Cells and transfection

Human kidney cell line HEK-293 and lung cancer cell line A549 were purchased from the Cell Bank of Type Culture Collection Committee of the Chinese Academy of Sciences. All cells were cultured in Dulbecco’s Modified Eagle Medium (DMEM high glucose, Gibco) containing 10% FBS (Biological Industries) and at 37 °C in 5% CO2 incubator. For transient overexpression assays, logarithmically growing cell lines were seeded into 6-well plates, then 3 ug of plasmids expressing SARS-CoV-2 sequence or vector were transfected with 5 μL jetPEI (Polyplus, USA) according to the manufacturer’s instructions. Six hours later, medium was refreshed and cells were harvested for further experiments 36 h after transfection. For drug treatment assays, fresh medium containing DMSO (Sigma, USA), 10 μM polydatin (MCE,USA) or 10 μM lonidamine (MCE,USA) was used after transfection.

### Protein isolation and western blot assay

Total cell protein was isolated with RIPA lysis buffer (50 mM Tris • HCl, pH 7.4, 0.1% SDS, 150 mM NaCl, 1 mM EDTA, 1 mM EGTA, 10 mM NaF) supplemented with protease inhibitor mixture (Roche, USA). After centrifugation at 13000 rpm for 30 min at 4°C, 20-30 μg protein was separated by 10% SDS-PAGE and transferred to PVDF membranes (Millipore, USA). After being blocked with 5% non-fat milk for 1 hour, membranes were incubated with the indicated primary antibodies overnight following the manufacturer’s instructions. Finally, membranes were incubated with horseradish peroxidase-conjugated secondary antibodies and protein expression was detected using an enhanced chemiluminescence (ECL) system (Thermo pierce, USA). The antibodies used in this study were the following: rabbit anti-RBD polyclonal antibody (Sino Biological, #40592-T62), mouse anti-Actin monoclonal antibody (ProteinTech, 60008-1-Ig), mouse anti-his tag monoclonal antibody (Abmart, M20001S), mouse anti-DDK/Flag tag monoclonal antibody (Abmart, M20008L), HRP Goat Anti-Rabbit IgG (ZSBIO, ZB-5301), HRP Goat anti-mouse IgG (ZSBIO, ZB-5305).

### Immunofluorescence assay

A549 cells were seeded on glass coverslips in 12-well plates and transfected with SARS-CoV-2-derived plasmids or control vector according to the method described above. Then cells were washed with cold PBS buffer and fixed in 4% paraformaldehyde. After treatment with 1% triton-100 for 15 min and blocking with unimmunized goat serum for 1 h, cells were incubated with the anti-his tag antibody overnight at 4°C, and then detected with donkey anti-mouse secondary antibody conjugated with alexa fluor plus 488nm (Invitrogen, USA). Cell nuclei were stained with DAPI staining buffer (Invitrogen, USA) and images were collected with a fluorescence microscope (Olympus, Japan).

### Virus replication assay

A type-B recombinant human adenoviral vector Ad11-5ETel-GFP with adenovirus E1A enhancer and human telomerase reverse transcriptase (TERT) promoter was constructed according our previous study(*34*). In order to evaluate the effect of polydatin and lonidamine on virus replication, 2×10^5^ A549 cells were seeded into 12-wells plates and infected with Ad11 virus (MOI=0.5) in serum-free medium. 2 hours later, medium was refreshed with 2% fetal calf serum containing DMSO or 10 μM mitochondria-targeted drugs. All cells and medium were harvested at 72 hours after infection, then frozen and thawed three times. The amounts of virus in cell lysates were measured with HEK-293 cell as previously described(*35*).

### Bioinformatics analysis

The differences in codon usage preference between SARS-CoV-2-derived protein coding sequences and human were analyzed using GCUA tools. The codon usage table of mitochondrial and other viruses used in this study were downloaded from the Codon Usage Database (http://www.kazusa.or.jp/codon/). Public gene expression data for SARS-CoV-2-infected cells (GSE147507) were obtained from the GEO database, and the differentially expressed genes and heatmap of selected genes were analyzed with R studio. Potential TFs on the promoter (upstream 1000bp) of TRL-TAA-4-1 were predicted with JASPAR software (relative profile score threshold was 80%) and Venny 2.1 software was used to performed the gene overlap analysis. The signalling pathway and functional enrichment analysis of related genes were performed with Metascape. The distribution of genes including ACE2, TMPRSS2, ATF2 and overlapped TFs in different cells of human lung tissue was analyzed with online tools in the single cell portal database (https://singlecell.broadinstitute.org/single_cell/study/SCP814/human-lung-hiv-tb-co-infection-ace2-cells).

## Statistical analysis

All statistical graphs and difference analysis between groups were evaluated with Graphpad Prism 7 using Student’s t-test (unpaired, two-tailed). p<0.05 was considered to indicate a significant difference.

## Conflict of interest

The authors declare no conflict of interest.

## Acknowledgement

This study was supported by grants from the National Natural Science Foundation of China (81702383, U1704282) and National Key Technologies R & D Program of China (2016YFE0200800).

## Notes

### Competing Interest Statement

The authors have declared no competing interest.

## REFERENCES AND NOTES

1. C. Shen et al., Treatment of 5 Critically Ill Patients With COVID-19 With Convalescent Plasma. Jama 323, 1582 (Apr 28, 2020).

2. K. Dhama et al., Coronavirus Disease 2019-COVID-19. Clinical microbiology reviews 33, (Sep 16, 2020).

3. D. E. Gordon et al., A SARS-CoV-2 protein interaction map reveals targets for drug repurposing. Nature 583, 459 (Jul, 2020).

4. D. Blanco-Melo et al., Imbalanced Host Response to SARS-CoV-2 Drives Development of COVID-19. Cell 181, 1036 (May 28, 2020).

5. D. Bojkova et al., Proteomics of SARS-CoV-2-infected host cells reveals therapy targets. Nature 583, 469 (Jul, 2020).

6. M. Bouhaddou et al., The Global Phosphorylation Landscape of SARS-CoV-2 Infection. Cell 182, 685 (Aug 6, 2020).

7. J. B. Plotkin, G. Kudla, Synonymous but not the same: the causes and consequences of codon bias. Nature reviews. Genetics 12, 32 (Jan, 2011).

8. C. H. Yu et al., Codon Usage Influences the Local Rate of Translation Elongation to Regulate Co-translational Protein Folding. Molecular cell 59, 744 (Sep 3, 2015).

9. L. Tian, X. Shen, R. W. Murphy, Y. Shen, The adaptation of codon usage of +ssRNA viruses to their hosts. Infection, genetics and evolution: journal of molecular epidemiology and evolutionary genetics in infectious diseases 63, 175 (Sep, 2018).

10. F. Valdez et al., Schlafen 11 Restricts Flavivirus Replication. Journal of virology 93, (Aug 1, 2019).

11. A. Nunes et al., Emerging Roles of tRNAs in RNA Virus Infections. Trends in biochemical sciences 45, 794 (Sep, 2020).

12. N. Zhu et al., A Novel Coronavirus from Patients with Pneumonia in China, 2019. The New England journal of medicine 382, 727 (Feb 20, 2020).

13. D. Kim et al., The Architecture of SARS-CoV-2 Transcriptome. Cell 181, 914 (May 14, 2020).

14. Zhenguo Cheng et al. (2020).

15. K. M. Lee, C. J. Chen, S. R. Shih, Regulation Mechanisms of Viral IRES-Driven Translation. Trends in microbiology 25, 546 (Jul, 2017).

16. L. C. de Oliveira et al., The Host Factor Early Growth Response Gene (EGR-1) Regulates Vaccinia virus Infectivity during Infection of Starved Mouse Cells. Viruses 10, (Mar 21, 2018).

17. J. Buehler et al., Host signaling and EGR1 transcriptional control of human cytomegalovirus replication and latency. PLoS pathogens 15, e1008037 (Nov, 2019).

18. F. A. Vieira Braga et al., A cellular census of human lungs identifies novel cell states in health and in asthma. Nature medicine 25, 1153 (Jul, 2019).

19. N. Zhu et al., Morphogenesis and cytopathic effect of SARS-CoV-2 infection in human airway epithelial cells. Nature communications 11, 3910 (Aug 6, 2020).

20. M. Hoffmann et al., SARS-CoV-2 Cell Entry Depends on ACE2 and TMPRSS2 and Is Blocked by a Clinically Proven Protease Inhibitor. Cell 181, 271 (Apr 16, 2020).

21. F. Boos, M. Wollin, J. M. Herrmann, Methionine on the rise: how mitochondria changed their codon usage. The EMBO journal 35, 2066 (Oct 4, 2016).

22. P. H. Willems, R. Rossignol, C. E. Dieteren, M. P. Murphy, W. J. Koopman, Redox Homeostasis and Mitochondrial Dynamics. Cell metabolism 22, 207 (Aug 4, 2015).

23. S. Chakraborty et al., Codon usage and expression level of human mitochondrial 13 protein coding genes across six continents. Mitochondrion 42, 64 (Sep, 2018).

24. K. E. Wu, F. M. Fazal, K. R. Parker, J. Zou, H. Y. Chang, RNA-GPS Predicts SARS-CoV-2 RNA Residency to Host Mitochondria and Nucleolus. Cell systems 11, 102 (Jul 22, 2020).

25. E. Lau, Z. A. Ronai, ATF2 - at the crossroad of nuclear and cytosolic functions. J Cell Sci 125, 2815 (Jun 15, 2012).

26. M. A. Nelo-Bazan et al., Early growth response 1 (EGR-1) is a transcriptional regulator of mitochondrial carrier homolog 1 (MTCH 1)/presenilin 1-associated protein (PSAP). Gene 578, 52 (Mar 1, 2016).

27. M. L. DeDiego et al., Severe acute respiratory syndrome coronavirus envelope protein regulates cell stress response and apoptosis. PLoS pathogens 7, e1002315 (Oct, 2011).

28. B. M. Ellen ter et al., Resveratrol And Pterostilbene Potently Inhibit SARS-CoV-2 Infection In Vitro. BioRxiv, (2020).

29. D. Cervantes-Madrid, Y. Romero, A. Duenas-Gonzalez, Reviving Lonidamine and 6-Diazo-5-oxo-L-norleucine to Be Used in Combination for Metabolic Cancer Therapy. Biomed Res Int 2015, 690492 (2015).

30. A. P. West et al., Mitochondrial DNA stress primes the antiviral innate immune response. Nature 520, 553 (Apr 23, 2015).

31. K. K. Singh, G. Chaubey, J. Y. Chen, P. Suravajhala, Decoding SARS-CoV-2 hijacking of host mitochondria in COVID-19 pathogenesis. Am J Physiol Cell Physiol 319, C258 (Aug 1, 2020).

32. S. K. Thaker, J. Ch’ng, H. R. Christofk, Viral hijacking of cellular metabolism. BMC biology 17, 59 (Jul 18, 2019).

33. F. Liu et al., A Novel Pak1/ATF2/miR-132 Signaling Axis Is Involved in the Hematogenous Metastasis of Gastric Cancer Cells. Molecular therapy. Nucleic acids 8, 370 (Sep 15, 2017).

34. H. H. Wong et al., Modification of the early gene enhancer-promoter improves the oncolytic potency of adenovirus 11. Molecular therapy: the journal of the American Society of Gene Therapy 20, 306 (Feb, 2012).

35. Y. Wang et al., E3 gene manipulations affect oncolytic adenovirus activity in immunocompetent tumor models. Nature biotechnology 21, 1328 (Nov, 2003).

