## Supplemental Figure1 for "A novel viral protein translation mechanism reveals mitochondria as a target for antiviral drug development"

Figures.S1

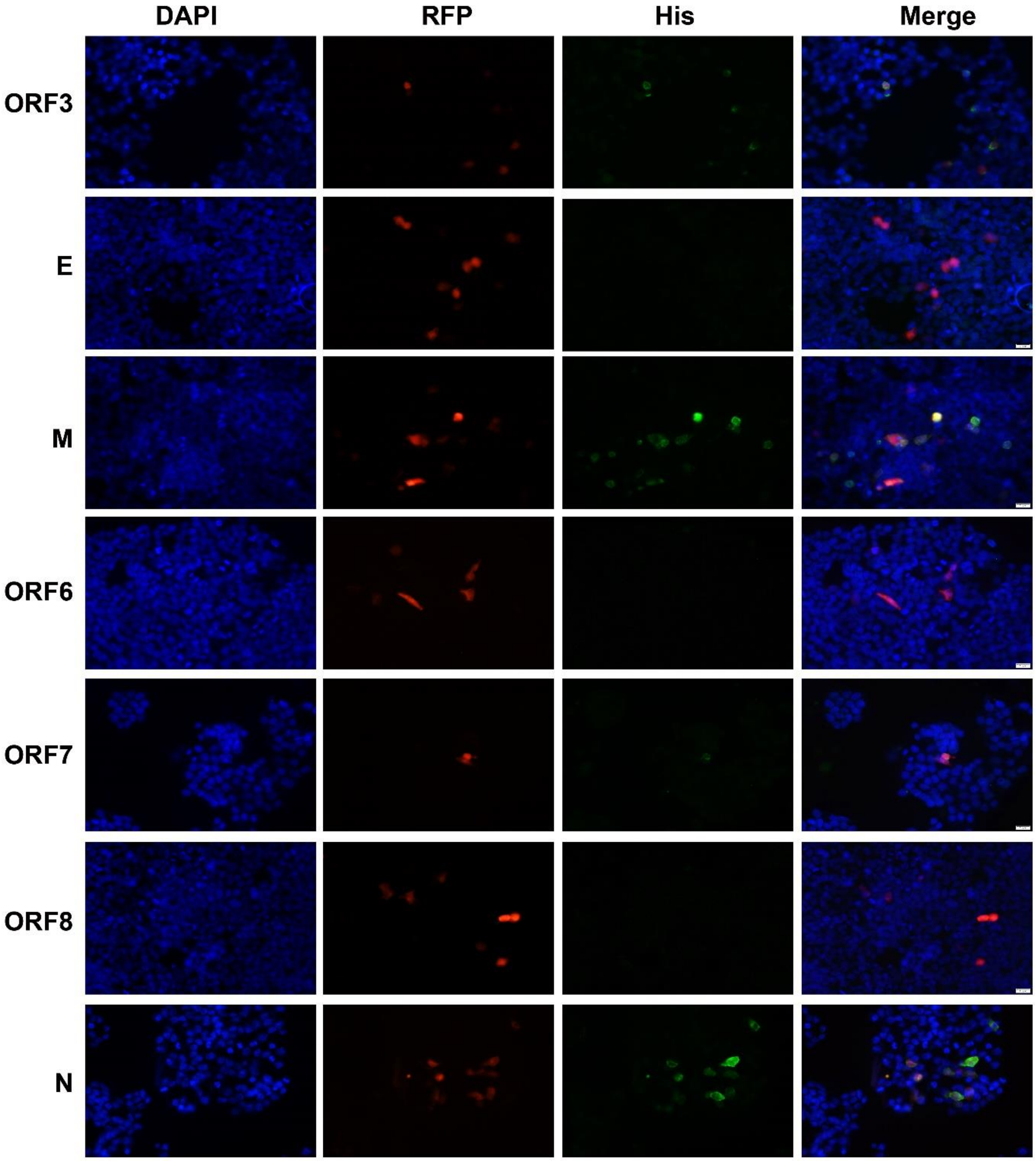

Fig S1. Protein expression of SARS-CoV-2 derived proteins including ORFs , E, M and N in human lung cancer cell line A549 cells after transfection with plasmids expressing different virus genes  by confocal microscopy.  All plasmids were constructed with IRES-RFP sequence.
